# Serotonin depletion amplifies distinct human social emotions as a function of individual differences in personality

**DOI:** 10.1101/789669

**Authors:** Jonathan W Kanen, Fréderique E Arntz, Robyn Yellowlees, Rudolf N Cardinal, Annabel Price, David M Christmas, Annemieke M Apergis-Schoute, Barbara J Sahakian, Trevor W Robbins

## Abstract

Serotonin is involved in a wide range of mental capacities essential for navigating the social world, including emotion and impulse control. Much recent work on serotonin and social functioning has focused on decision-making. Here we investigated the influence of serotonin on human emotional reactions to social conflict. We used a novel computerised task that required mentally simulating social situations involving unjust harm and found that depleting the serotonin precursor tryptophan – in a double-blind randomised placebo-controlled design – enhanced emotional responses to the scenarios in a large sample of healthy volunteers (n = 73), and interacted with individual differences in trait personality to produce distinctive human emotions. Whereas guilt was preferentially elevated in highly empathic participants, annoyance was potentiated in those high in trait psychopathy, with medium to large effect sizes. Our findings show how individual differences in personality, when combined with fluctuations of serotonin, may produce diverse emotional phenotypes. This has implications for understanding vulnerability to psychopathology, determining who may be more sensitive to serotonin-modulating treatments, and casts new light on the functions of serotonin in emotional processing.

## Introduction

A unified function for serotonin (5-hydroxytryptamine; 5-HT) has, perhaps unsurprisingly, proven to be elusive. It is hypothesised to have a role in many psychiatric disorders ^1^, and is implicated in a wide range of mental functions including aversive processing, impulse control, and social behaviour ^2^. These domains can be viewed under a unified framework by considering how serotonin impacts Pavlovian (stimulus-outcome, including emotional) and instrumental (stimulus-response-outcome; operant) processes that underlie both social and non-social functions ^3^. Much recent work on the relationship between serotonin, aversive processing, and human social behaviour has focused on instrumental action in the context of behavioural economic games and moral dilemmas ^4–6^. Here we examined emotional reactions to social scenarios depicting unjust harm.

Studies of healthy volunteers have primarily employed two techniques to investigate serotonin function: acute tryptophan depletion (ATD), a dietary technique that temporarily lowers brain serotonin levels by depleting its biosynthetic precursor, tryptophan ^7–11^, and treatment with single doses of selective serotonin reuptake inhibitors (SSRIs). SSRIs are generally assumed to increase extracellular serotonin; however, it should be noted that single rather than chronic doses can paradoxically decrease serotonin in projection areas ^2,12^. ATD studies have revealed disinhibition of retaliatory behaviour in the face of perceived injustice ^4^, modelled using the Ultimatum Game (UG), whilst single dose SSRI has had pro-social effects ^5^.

Morally relevant social emotions, meanwhile, lie at the interface between moral standards and socially appropriate behaviour ^13^. Moral standards prohibit behaviours that are likely to have negative consequences for the well-being of others. Emotions are in part Pavlovian in nature, and whilst ATD has been shown to modulate Pavlovian processes in non-social domains ^14,15^, here we tested the influence of ATD on emotion in a social context. We asked whether ATD would enhance morally relevant negative emotions evoked by social scenarios in which a person was unjustly harmed: these served as Pavlovian cues. Using a novel task, we prompted participants to reflect on the situations and specifically assessed emotions involving annoyance, guilt, shame, and feeling “bad”. Reporting one’s emotional reactions after mentally simulating hypothetical social scenarios implicitly calls on autobiographical memories, which should result in a diversity of subjective experiences tied to personal qualities and expectations about social behaviour. We therefore also tested the influence of three personality traits – empathy, psychopathy, and impulsivity – on how serotonin would affect emotion.

Research on moral emotions has focused mostly on the self-conscious negative emotions guilt and shame ^13^. Guilt often relates to a negative appraisal of a specific behaviour, whereas shame tends to involve a negative evaluation of the self ^13^: “If only I hadn’t” as opposed to “If only I weren’t” ^16^. Whilst guilt is part of the diagnostic criteria for depression ^17^, proneness to shame most consistently relates to an array of psychiatric conditions, including symptoms of depression ^18–20^, anxiety ^19^, post-traumatic stress disorder ^21–24^, and eating disorders ^25,26^, as well as more specific symptoms such as low self-esteem ^27^, suicidal ideation ^28^, anger ^29,30^, and aggression ^30^. Importantly, guilt is thought to become maladaptive primarily when it is fused with shame ^13^.

Evidence from patients with damage to the ventromedial prefrontal cortex (vmPFC) and incarcerated individuals with psychopathy provides a plausible connection between moral emotions and social behaviour. The vmPFC is an area central to emotion regulation ^31^, with dense serotonergic innervation ^32^. Individuals with vmPFC damage also show increased retaliatory behaviour to unfairness on the UG ^33^, which mirrors the ATD findings ^4,6^. This is furthermore analogous to the UG results from psychopathic individuals ^34^, where vmPFC dysfunction is a feature ^35^, as is reduced guilt ^36^. Impaired moral behaviour following damage to the vmPFC has meanwhile been conceptualised as a manifestation of diminished guilt ^37^. The implication here is that guilt is related to inhibition of anti-social behaviour, as modelled by restraint in behavioural economic games like the UG. Studying the effects of ATD on guilt could therefore inform how moral emotion and behaviour are integrated.

Proneness to guilt consistently correlates with empathy, which refers to the ability to share the affective experiences of others ^13^. Empathy, whilst not a discrete emotion, is a morally relevant emotional capacity ^13^ and trait ^38^. Guilt appears to foster reparative action, promote empathy ^13^, and increase altruistic acts ^39^. Elevated empathy, moreover, has been correlated with severity of depression, and proposed as a risk factor for its development ^40^. Importantly, empathy is classically absent in psychopathy ^41^.

We were especially interested in empathy because moral decision-making in individuals high in trait empathy has been shown to be particularly sensitive to manipulations of serotonin ^5^. Furthermore, the serotonin 2A (5-HT2A) receptor agonists lysergic acid diethylamide (LSD), psilocybin, and 3,4-methylenedioxymethamphetamine (MDMA), have all been shown to enhance empathy ^42–44^. To extend these findings we therefore tested the hypothesis that ATD would interact with trait empathy to amplify morally relevant social emotion. Given the consistent correlations between guilt-proneness and empathy ^13^, we predicted guilt would be the most likely emotion to be affected.

Conversely, psychopathy is characterised by emotional dysfunction reflected in reduced guilt and empathy ^45^, and an increased risk for aggression. Aggression can be either goal-directed (e.g. a premeditated crime), or reactive: an explosive, impulsive response to frustration ^36^. Many psychiatric conditions increase the risk for reactive aggression, however psychopathy is unique in that there is an increased risk for both reactive and goal directed aggression ^36^. Aggression is in turn traditionally associated with low serotonin ^46^, including evidence from studies of violent offenders ^47^. We explored whether psychopathic traits are likewise related to morally relevant emotions, and whether ATD modulates this relationship. We predicted that psychopathic traits might have the most pronounced effect on feeling annoyed, which invokes the notion of frustration.

Whilst some, but not all, aggression can be impulsive, aggression and impulsivity are distinct. Indeed, discrete serotonergic circuits modulate aggressive versus impulsive behaviour in mice ^48^. ATD can induce “waiting impulsivity” (diminished action restraint whilst waiting for a reward) and “impulsive choice” (accepting small immediate rewards over larger delayed ones) in healthy individuals ^6,49^. More impulsive choice has been correlated with increased aggressive impulses to perceived injustice on the UG, which was heightened further by ATD ^6^. We therefore asked whether trait impulsivity on the Barratt Impulsiveness Scale ^50^ was related to increased annoyance following ATD.

To test our hypotheses, we employed ATD in a double-blind, randomised, placebo-controlled, between-groups design, in healthy volunteers. We predicted that ATD would enhance negative emotion overall, and that individual differences in empathy, psychopathy, and impulsivity would influence how ATD modulated the profile of emotion. In line with the traditional disconnection between psychopathy and empathy, we predicted that there would be dissociation between how trait empathy and psychopathy interact with neurochemical status to modulate annoyance, guilt, shame, and feeling bad. Given the established connection between serotonin and impulsivity ^49,51,52^, and retaliatory behaviour ^6^, we also hypothesised that high trait impulsivity would be related to increased feelings of annoyance, which would be further potentiated by ATD in these individuals.

## Materials and Methods

### Participants

Seventy-three healthy participants (39 males, mean age 24.6) completed the experiment. Participants were medically healthy and screened to be free from any psychiatric disorders, using the Mini-International Neuropsychiatric Interview ^53^. Individuals who reported, during screening, having a first-degree relative (parent or sibling) with a psychiatric disorder were also excluded. Other exclusion criteria included neurological disorders, current use of any regular medication excluding contraceptive pills, and past use of neurological or psychiatric medication. Further exclusion criteria and measures to ensure a matching of the groups are reported in the Results and Supplementary Material. Participants gave informed consent before the start of the study and were paid for their participation.

### General Procedure

The protocol was approved by the Cambridge Central Research Ethics Committee (16/EE/0101), and the study took place at the National Institute for Health Research / Wellcome Trust Clinical Research Facility at Addenbrooke’s Hospital in Cambridge, England. Participants arrived in the morning of the study day having fasted for at least 9 hours beforehand and completed a baseline 16-item visual analogue scale (VAS) to assess mood and other feelings including alertness. The VAS was also completed during the middle and end of the day. Participants then gave a baseline blood sample, and ingested either a placebo or tryptophan depletion drink. Simple randomization was employed. In the afternoon, participants gave a second blood sample, approximately 4.5 hours after ingesting the drink, and completed the Moral Emotions task (described below). Other tasks that will be reported elsewhere preceded the Moral Emotions task in the following order and examined: behavioural attributions ^54^, probabilistic reversal learning ^55^, theory of mind ^56^, Pavlovian conditioning ^57^, avoidance learning ^58^, and deterministic reversal learning. The administration of multiple tasks within a single ATD study is a common approach taken by several studies cited within this manuscript ^4,6,14,49,59^. Participants additionally attended a short afternoon session the day before, with no pharmacological manipulation, where they completed baseline questionnaires and three unrelated tasks, which will not be reported here. The present laboratory experiment was conducted once.

### Acute Tryptophan Depletion

Tryptophan is the amino acid precursor necessary to synthesize brain serotonin. Acute tryptophan depletion (ATD) is a widely used dietary manipulation, which results in a rapid decrease in the synthesis and release of serotonin ^7–11^. Participants were randomly assigned to receive either ATD or a placebo condition, in a double-blind, between-groups design. The depletion group received a drink that contained a balance of all the essential amino acids except for tryptophan. The placebo group received the same drink except that it included tryptophan. Blood plasma samples were collected to verify depletion and analysed using high performance liquid chromatography (HPLC).

### Moral Emotions Task

We employed a novel task, part of the EMOTICOM neuropsychological testing battery ^56,60^, to measure feelings of guilt, shame, annoyance, and feeling “bad”. Using a touchscreen computer, participants were presented with cartoons of social scenarios – Pavlovian cues – in which someone was unjustly harmed, either intentionally or unintentionally. We then interrogated emotional reactions to these scenarios by asking participants to reflect, and report how they would feel if they were the victim or agent of harm. An example of one trial is depicted in Figure 1. There were 28 randomised trials, composed of 14 different cartoons, which were each presented twice – once where participants were prompted to identify as the victim, and once where they were asked to identify as the agent. The specific instruction was, “If this was you, please indicate below how you would feel by touching the line.” The four different emotions were measured using four unnumbered touchscreen scales, with seven rungs to choose from, where the first rung was labelled “not at all”, scored as 1, and the seventh labelled “extremely”, scored as 7. Half of the scenarios involved an intentional harm. In the other half, the harm committed was accidental. The task was self-paced.

**Figure 1.**
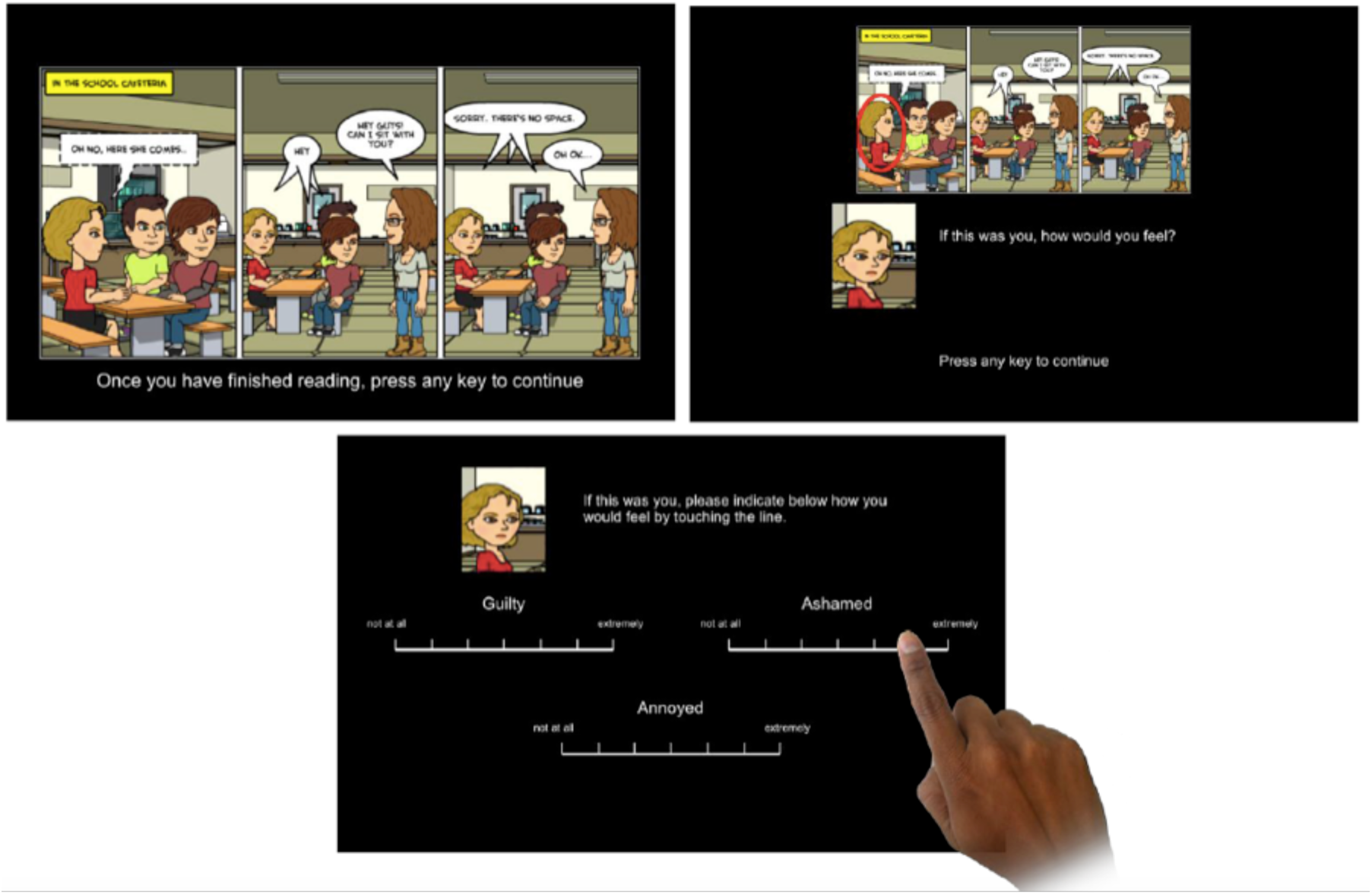
Moral emotions task schematic. Three example slides of a trial are shown. Feeling “bad” was assessed with a rating scale on a fourth slide.

### Statistics

Sample size was chosen based on a recent analogous study that also employed the EMOTICOM neuropsychological testing battery, in healthy volunteers, and using a serotonergic manipulation ^61^. Data were analysed using MATLAB (MathWorks) and SPSS (IBM). MATLAB code is available as supplementary material. All statistical tests were two-sided. Homogeneity of variance in t-tests was verified with Levene’s test, and degrees of freedom were adjusted when this assumption was violated. The Greenhouse-Geisser correction was used where applicable, in designs with within-subjects factors, to correct for violation of the sphericity assumption as determined by Mauchly’s test. Adjustments for multiple comparisons were not deemed necessary.

## Results

Seventy-three participants completed the study: thirty-seven underwent depletion (20 males), while the remaining 36 received placebo (19 males).

### Effects of ATD on mood ratings

Mood ratings were unaffected by ATD. We collected rating data from 65 participants (n = 33 depletion) on how happy or sad they were feeling before the task, after depletion had taken effect, and these ratings did not differ from those of participants in the placebo group (t_(63)_ = −0.727, p =. 47). There were also no baseline differences in depressive symptoms (t_(71)_ = −1.258, p =. 212) using the Beck Depression Inventory-II ^62^.

### How serotonin depletion modulates emotional ratings overall

We tested whether serotonin depletion potentiated emotions overall, irrespective of individual differences. To do this we performed repeated measures analysis of variance (ANOVA) incorporating all four emotions measured. Serotonin status (ATD versus placebo) was the between-subjects factor, and the within-subjects factors were emotion (annoyance, guilt, shame, and feeling bad), agency (agent or victim of harm), and intentionality (intentional or unintentional harm). Indeed, serotonin depletion potentiated emotion overall (F_(1,71)_ = 5.959, p =. 017, η_p_^2^ =. 077), shown in Table 1 and Supplementary Figure 1. There were highly significant main effects for emotion (F_(2,155)_ = 171.744, p = 7.4 × 10^−42^, η_p_^2^ =. 708), agency (F_(1,71)_ = 630.212, p = 4.9 × 10^−37^, η_p_^2^ =. 899), and intentionality (F_(1,71)_ = 11.799, p =. 001, η_p_^2^ =. 143), regardless of serotonergic status, which supports task validity and suggests individuals were attuned to the social context of the scenarios. There were no interactions between serotonin status and agency or intentionality (all p >. 05). Our core results in this study however came from analyses of how individual differences interacted with the serotonin-depleted state to modulate emotion, which now follow.

**Table 1.** Summary of results on personality traits. **↑↑**indicates a significant enhancement of the emotion by ATD at high levels of the personality trait shown; **+**indicates a significant positive relationship between the personality trait and the emotion that was not significantly modulated by ATD; **−** likewise indicates a significant negative relationship. Emotions are collapsed across agency and intentionality.

|  | Annoyance |  | Guilt |  | Shame |  | Bad |  |
| --- | --- | --- | --- | --- | --- | --- | --- | --- |
|  | Placebo | ATD | Placebo | ATD | Placebo | ATD | Placebo | ATD |
| Impulsivity |  |  |  |  |  |  | - |  |
| Psychopathy | | $\uparrow\uparrow$ | | | | | | |
| Empathy | | | | $\uparrow\uparrow$ | + | + | | |

### Correlations between trait measures

First, we ensured that our trait measures of interest were not correlated with one another. Scores on the impulsivity and psychopathy scales were not correlated (r_(73)_ = -.113, p =. 342). Empathy scores were likewise not correlated with impulsivity (r_(73)_ =. 094, p =. 427) or psychopathy (r_(73)_ =. 015, p =. 897).

### How trait empathy modulates emotional effects of serotonin depletion

Given prior evidence that single dose SSRI had a more pronounced effect on social behaviour in highly empathic participants ^5^, a central question in our study was whether the serotonin-depleted state and empathic trait interacted to influence emotion. Indeed, we found that the self-conscious emotion guilt was more sensitive to serotonin depletion in individuals with high trait empathy. We ensured self-reported empathy at baseline, using the Interpersonal Reactivity Index (IRI) ^38^, was matched between placebo and depletion (t_(63)_ = 0.442, p =. 660; Levene’s test F_(71)_ = 4.569, p =. 036). We analysed emotional ratings via analysis of covariance (ANCOVA), using empathy and serotonin status (ATD versus placebo) as factors in a between-subjects interaction term, controlling for main effects, and agency (agent or victim) and intent (intentional or unintentional) as within-subjects factors. For guilt ratings, this revealed a significant four-way serotonin × empathy × agency × intentionality interaction (F_(1,69)_ = 5.596, p =. 021, η_p_^2^ =. 075). Guilt ratings were significantly higher in more empathic individuals following serotonin depletion (r_(37)_ =. 385, p =. 019), and not under placebo (r_(36)_ =. 265, p =. 118), irrespective of agency or intentionality, seen in Figure 2a and Table 1. Follow-up tests showed that the agency and intentionality effect was driven by a highly significant relationship between empathy and guilt, when imagining inflicting (being the agent of) harm unintentionally (r_(37)_ =. 43, p =. 008). There was additionally a three-way interaction between agency, intentionality, and group (F_(1,69)_ = 4.765, p =. 032, η_p_^2^ =. 065). Critically, however, this three-way interaction disappeared when empathy was not included as a factor in the (otherwise identical) model (F_(1,71)_ =. 608, p =. 438, η_p_^2^ =. 008). We were also able to reproduce this core result on guilt, using a quartiles approach whereby individuals with an empathy score in the top 25% were deemed “high empathy” and the bottom 25% “low empathy”. ANCOVA with empathy (high versus low) and serotonin status (ATD versus placebo) as factors in a between-subjects interaction term, controlling for main effects, and agency (agent or victim) and intent (intentional or unintentional) as within-subjects factors, again revealed a significant four-way serotonin empathy agency intentionality interaction effect on guilt (F_(1,36)_ = 7.028, p =. 012, η_p_^2^ =. 163). Whilst the ANCOVA approach, as above, did not yield a significant interaction of serotonin and empathy on shame (F <. 42, p >. 05, η_p_^2^ <. 007 for all terms involving serotonin status), empathy and overall shame ratings were highly correlated in both the depletion (r_(37)_ =. 437, p =. 007) and placebo groups (r_(36)_ =. 425, p =. 01). These data for shame are shown in Figure 2b and Table 1. The ANCOVA on annoyance was not significant (F < 2, p >. 05, η_p_^2^ <. 03 for all terms involving serotonin status), as was the case for the ANCOVA on feeling “bad” (F < 1.2, p >. 05, η_p_^2^ =. 02 for all terms involving serotonin status). Guilt, therefore, was uniquely affected by the interaction between trait empathy and the serotonin-depleted state. Serotonin induced a distinct emotional profile in highly empathic individuals.

**Figure 2.**
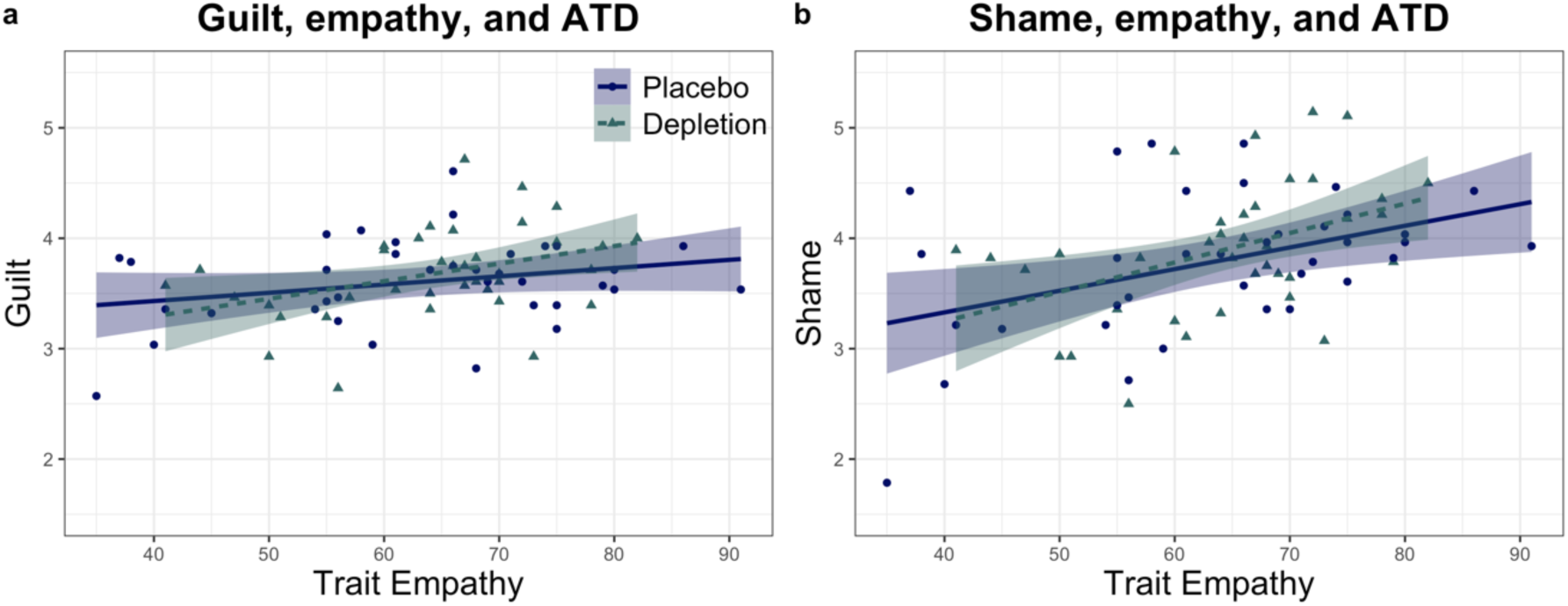
Effects of trait empathy on how serotonin depletion influences emotion. Shading indicates 1 standard error. Each point represents the average emotion ratings for each individual, collapsed across agency and intentionality. **a)** The highly empathic reported more guilt following depletion relative to when on placebo. **b)** Shame was significantly elevated in individuals high in trait empathy on both placebo and depletion.

### How trait psychopathy modulates emotional effects of serotonin depletion

Trait psychopathy also interacted with the serotonin-depleted state to modulate emotion. We ensured psychopathic traits assessed with the Levenson Self-Report Psychopathy Scale 63 at baseline were matched in the placebo and depletion groups (t_(71)_ = 1.132, p =. 261). ANCOVA with serotonin status (ATD versus placebo) and psychopathy as factors in a between-subjects interaction term, controlling for main effects, and agency and intentionality as within-subjects factors revealed a significant serotonin × psychopathy × intentionality three-way interaction for feelings of annoyance (F_(1,69)_ = 7.172, p =. 009, η_p_^2^ =. 094). With increasing trait psychopathy, individuals felt even more annoyed following serotonin depletion, seen in Figure 3a and Table 1. Intentionality significantly interacted with psychopathy to influence annoyance under placebo (F_(1,34)_ = 5.163, p =. 03, η_p_^2^ =. 132) and not following depletion (F_(1,34)_ = 2.237, p =. 144, η_p_^2^ =. 06). In other words, the influence of psychopathy on annoyance depended on intentionality when on placebo, but on depletion those high in trait psychopathy were more annoyed regardless of intentionality. There was additionally a significant serotonin intentionality interaction (F_(1,69)_ = 7.161, p =. 009, η_p_^2^ =. 094). Critically, however, this two-way interaction disappeared when psychopathy was not included as a factor in the (otherwise identical) model (F_(1,71)_ =. 043, p =. 836, η_p_^2^ =. 001). Next, we assessed guilt using the same ANCOVA approach: there was no serotonin psychopathy interaction (F_(1,69)_ < 2.8, p >. 05, η_p_^2^ <. 04 for all terms involving serotonin status). There was no serotonin-by-psychopathy interaction for shame (F_(1,69)_ < 3.8, p >. 05, η_p_^2^ <. 055 for all terms involving serotonin status) nor for feeling bad (F_(1,69)_ < 2.1, p >. 05, η_p_^2^ <. 03 for all terms involving serotonin status).

**Figure 3.**
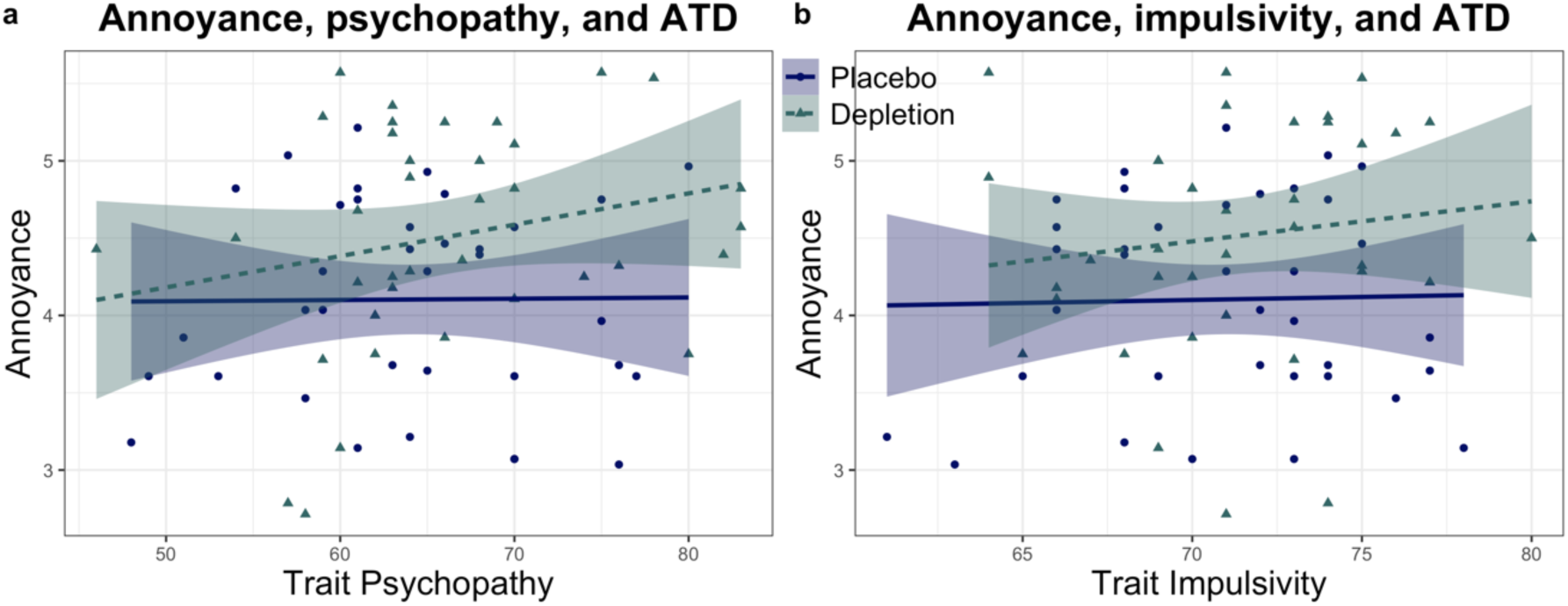
Effects of trait psychopathy and impulsivity on emotional effects of serotonin depletion. Shading indicates 1 standard error. Each point represents the average emotion ratings for each individual, collapsed across agency and intentionality. **a)** Annoyance was potentiated by serotonin depletion in high trait psychopathy. **b)** Trait impulsivity did not significantly enhance the effects of ATD on annoyance.

### Trait impulsivity and emotional effects of serotonin depletion

Trait impulsivity, measured at baseline with the Barratt Impulsiveness Scale ^50^, was also matched between groups, and did not interact with the serotonin-depleted state to modulate emotion. Data on impulsivity are summarised in Table 1. First, we assessed feelings of annoyance using ANCOVA with serotonin status (ATD versus placebo) and impulsivity as factors in a between-subjects interaction term, controlling for main effects, and agency (agent or victim) and intent (intentional or unintentional) as within-subjects factors: there was no interaction between serotonin and impulsivity (F_(1,69)_ < 1.7, p >. 05, η_p_^2^ <. 025 for all terms involving serotonin status). These data are shown in Figure 3b. The same was true for all terms involving serotonin status in the ANCOVAs on guilt (F_(1,69)_ <. 7, p >. 05, η_p_^2^ <. 01), shame (F_(1,69)_ <. 4, p >. 05, η_p_^2^ <. 01) and feeling bad (F_(1,69)_ < 2.6, p >. 05, η_p_^2^ <. 04).

### Principal Components Analysis

We also explored whether there was any structure underlying the task measurements that was not detected by our prior analyses. To do this, we used principal components analysis (PCA) on the 16 outcome variables from the task (see Supplementary for methods; factor loadings are shown in Supplementary Table 1). The validity of this PCA was confirmed by comparing the number of components, and the variables that clustered together, to a PCA on the same task in the original larger dataset from a non-pharmacological study of 186 healthy participants 56. We then interpreted how the task measurements from our experiment clustered into the four principal components. Component 1 centred on annoyance with others for having done harm to oneself – in other words, outward frustration. The predominant theme of component 2 was inward frustration, or annoyance with oneself for having harmed another. Components 3 and 4 centred on the self-conscious negative emotions guilt and shame. Component 3 captured these emotions when the participant was the agent, component 4 when the participant was the victim of harm. We then used the estimated factor scores for each individual to assess how serotonin depletion modulated the constructs captured by the four components. ANOVA with serotonin status (ATD versus placebo) as a between-subjects factor, and the four components as levels of a within-subjects factor, revealed a significant serotonin-by-component interaction (F_(3,213)_ = 3.165, p =. 025, η_p_^2^ =. 043). There was no main effect of serotonin depletion (F_(1,71)_ = 2.187, p =. 144, η_p_^2^ =. 030). Follow up t-tests revealed the values of component 2 (t_(71)_ = 2.124, p =. 037) and component 4 (t_(71)_ = 2.290, p =.025) were each significantly greater following depletion relative to placebo, as seen in Supplementary Figure 2. Inward frustration, or annoyance, for having harmed another – captured by component 2 – was potentiated by serotonin depletion. When the victim of harm in the task, self-conscious negative emotion was also exacerbated by serotonin depletion (component 4).

### Blood Analysis

Blood results are shown in Figure 4. The ratio of tryptophan to large neutral amino acids (TRP:LNAAs; tryptophan to valine, methionine, isoleucine, leucine, tyrosine, and phenylalanine) was calculated, as this is thought to be most reflective of the extent of brain serotonin depletion ^64^. We then performed a t-test on the change in the TRP:LNAA ratio between samples taken at baseline and approximately 4.5 hours following administration of the mixture. Plasma levels were unavailable for two participants: one due to a staff processing error, and one due to difficulty with venepuncture. We achieved a robust depletion of tryptophan (t_(60)_ = −19.163, p = 3.01 × 10^−27^).

**Figure 4.**
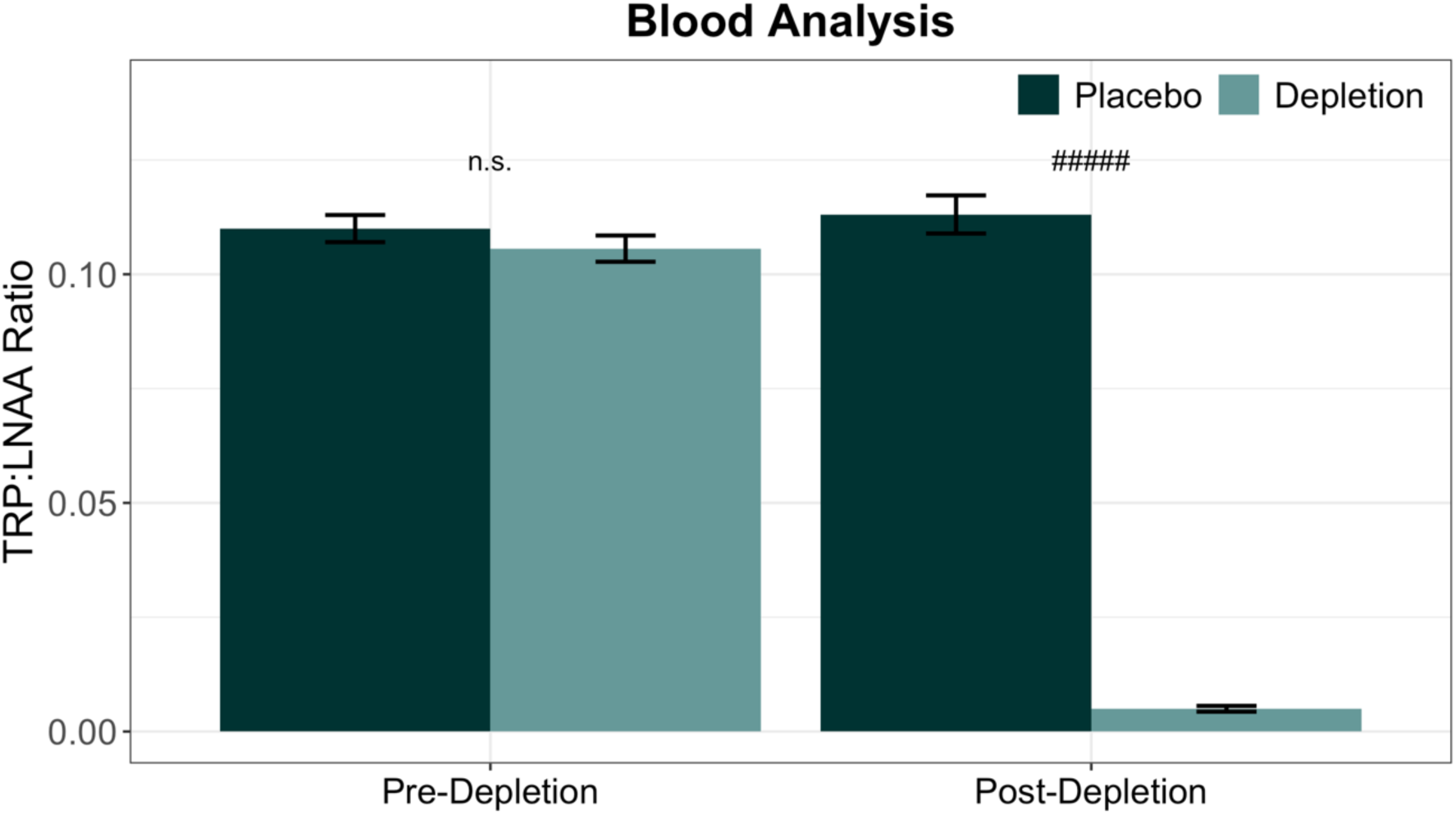
Blood analysis results. Error bars represent 1 standard error. Significance at p < 5 × 10^−27^ is denoted by #####. n.s. denotes not significant.

### Correlation analyse

Extent of depletion was not correlated with any particular emotion, reinforcing the importance of accounting for traits to uncover effects of ATD on social emotion. Change in TRP:LNAA ratio was not correlated with annoyance (r = -.208, p =. 081), guilt (r = -.121, p =. 314), shame (r = -.057, p =. 636), or feeling bad (r = -.088, p =. 464). There were correlations between emotion ratings: collapsed across serotonergic status, annoyance, guilt, and shame were all positively correlated with one another (p <. 05), consistent with the nonspecific enhancement of emotion in the absence of considering traits. Guilt and shame were both negatively correlated with feeling bad (p <. 05). These correlations, with exact p-values and correlation coefficients, are presented in Supplementary Table 2.

### Summary of results

Results are summarised in Table 1. Serotonin depletion enhanced emotion overall. Examining individual trait differences revealed a deeper story. Guilt was significantly enhanced by serotonin depletion in highly empathic participants, which was a distinctive emotional profile. This was driven by guilt when the agent of an unintentional harm. Trait empathy, furthermore, was highly correlated with shame ratings in both the placebo and depletion groups. Greater trait psychopathy following serotonin depletion, meanwhile, was associated with enhancement of annoyance. PCA revealed a component where guilt and shame clustered together when the victim of harm, which was potentiated by ATD.

## Discussion

Using a novel test that cued autobiographical memories, we showed that serotonin depletion heightened emotional reactions when mentally simulating social scenarios involving unjust harm. Whilst emotion was enhanced non-specifically at the group level, harnessing baseline individual differences revealed that personality traits play a critical role in shaping which distinctive types of emotions are affected by serotonin depletion. A key result was that individuals high in trait empathy showed a distinct profile of enhanced guilt following serotonin depletion. This contrasted with how variation in trait psychopathy (classically associated with lack of empathy and guilt) influenced the relationship between serotonin depletion and emotional reactions: only annoyance was potentiated. This dissociation, in other words, mirrored the antithetical nature of empathy and psychopathy. Previous studies have shown that traits can influence the magnitude of effects of serotonin manipulations on social behaviour ^5,6^; we now show that traits modulated the quality, as well as the magnitude, of social emotion following serotonin depletion. Traits contribute to an individual’s model of the world, and therefore shape prior expectations about social interactions: we propose the influence of traits on how serotonin modulated emotions can be thought of as constituting biological “priors”.

Emotions prepare the body for action or inaction ^65^. Our findings on the serotonergic modulation of social emotion converge with the literature on social decision-making, and we propose that our results represent a Pavlovian influence that can shape social behaviour ^3^. The deployment of empathy has additionally been described as having a Pavlovian character, which can shape behaviour triggered by cues signalling harm ^3^. Whilst we did not measure behaviour, there are some intriguing parallels between our findings on emotional reactions to unjust harm, and studies of retaliatory behaviour to unfairness, as assessed using the Ultimatum Game ^4,6^ which we highlight below. In the UG, individuals must split a sum of money with another, and are given the opportunity to reject unfair offers, in which case neither player gets paid ^4,6^. Indeed, the UG has been studied under serotonin depletion and in relation to empathy and psychopathy.

Empathy was of central importance to our analysis. This was motivated by the observation that social behaviour in highly empathic participants was especially sensitive to single dose SSRI administration: these individuals showed the greatest reduction in retaliatory behaviour to unfairness ^5^. Critically, we found that individuals high in trait empathy had a distinct profile of enhanced emotion: guilt was amplified. Empathy and guilt are consistently correlated ^13^, and guilt may even promote empathy ^13^. Feelings of guilt have been associated with real-life altruistic acts ^39^. Guilt has been proposed to restrain antisocial behaviour as modelled by laboratory tests: a diminished sense of guilt is thought to contribute to dysfunctional social behaviour following vmPFC damage ^37^, and is also a feature of psychopathy ^36^. The present observation that increased guilt in the highly empathic under ATD was driven by instances of unintentionally inflicting harm in a social setting is entirely consistent with these accounts.

Shame, furthermore, was highly correlated with empathy. This was true in both the placebo and depletion conditions, which could have led to a ceiling effect, leaving less room for further enhancement of shame by ATD in the highly empathic, on top of an already elevated baseline under placebo. Guilt and shame, moreover, clustered together in the PCA when the victim of harm, and this component was enhanced by serotonin depletion regardless of personality traits. Guilt and shame are distinguishable yet can overlap ^13^, which is also true of their neural correlates ^66^. Whilst guilt and shame were correlated, the literature indicates that different events can be guilt-inducing or shame-inducing for different people (Tangney et al. 2007). That different types of emotion were enhanced in tandem at the group level, furthermore, does not imply the quality of these emotions cannot be distinguished by the person experiencing them.

Shame-free guilt is seen as possibly adaptive – for instance by promoting reparations – whilst proneness to shame is seen as a risk for, and is indeed associated with a wide range of psychopathology ^13^. Importantly, guilt is thought to become maladaptive primarily when it is fused with shame ^13^. Guilt overlaid with shame is most likely a source of rumination ^13^. The hippocampus has been reported to be involved in the experience of shame ^67^, and failure of hippocampal serotonin is suggested to contribute to rumination ^68^. SSRIs, meanwhile, improve hippocampal function in depression ^69^. Individuals with hippocampal damage, moreover, appear to show heightened reactive emotionality congruent with their behaviour in moral decision-making tasks, which is antithetical to the pattern seen with vmPFC lesions ^70^. While the relationship with our results is merely speculative, hippocampal dysfunction is a feature of numerous psychiatric conditions, as is social dysfunction: a recent framework proposes these two well established phenomena can be unified through the purported role of the hippocampus in organising social information (memories), via relational maps that support simulations of social outcomes ^71^.

Indeed, there are reported links between elevated empathy and depression ^40^. Individuals sensitive to distress in others may be more likely to experience personal distress, and this has been highlighted as a vulnerability factor for depression ^39,40^. At the same time, there is evidence for diminished deployment of theory of mind – non-affective perspective taking – in depression ^72^, and this combination raises the possibility that sensitivity to distress in oneself and others may become misattributed or inappropriately directed inward.

Whilst mood was unaffected in our study, consistent with the literature on ATD in healthy individuals ^73^, ATD can transiently reinstate low mood in depressed individuals successfully treated with an SSRI ^73^. By using trait measurements and a task that elicited emotions, however, we were able to uncover a pattern reminiscent of depression: more guilt in the highly empathic under serotonin depletion. Indeed, this task has already been used to detect possible latent vulnerabilities in a healthy population with trait paranoia ^60^. We propose that empathy, which produced a qualitatively unique emotional profile under ATD, may represent an important proxy for sensitivity to changes in serotonin.

Conversely, psychopathic individuals classically have impairments in guilt and empathy, and an increased risk for aggressive behaviour, especially following frustration ^36,45^. This is consonant with our results. We found that the emotional profile following serotonin depletion in healthy individuals high in psychopathic traits dissociated from what we observed in the highly empathic: annoyance was instead amplified following unjust harm. This result is furthermore in line with existing literature on social decision-making: clinically psychopathic individuals show an analogous pattern of behaviour on the UG 34 to the disinhibited aggressive impulses seen in ATD studies of healthy volunteers ^4^, that is also quantitatively similar to how individuals with vmPFC lesions behave on the UG ^34^. Critically, vmPFC damage is associated with impaired emotion regulation, and individuals with such lesions tend to exhibit anger and irritability particularly following frustration in their personal lives ^33^. Diminished structural and functional connectivity between the vmPFC and amygdala in clinically psychopathic individuals ^35^ is indeed thought to be a central mechanism underlying the condition. Interactions between these structures are furthermore sensitive to ATD in healthy individuals viewing facial signs of aggression ^59^. That serotonin depletion made participants high in trait psychopathy more annoyed by social injustice may be relevant for understanding how serotonin affects the emotional basis of retaliatory behaviour to unfairness ^4,6^. This view is supported by work showing that such behavioural reactions are associated with self-reported anger in healthy volunteers ^74^. Trait anger and psychopathy in violent offenders indeed appears to reflect 5-HT1B receptor levels ^75^, which moreover fits with the vast literature implicating serotonergic dysfunction in aggression ^46,51,76^.

The individual differences we observed in response to a challenge of brain serotonin are likely in part related to the relative contribution of the multiple serotonin subsystems in the brain. Importantly, preferential dysfunction in the median or dorsal raphe nuclei, which innervate, among other regions, the hippocampus and prefrontal cortex, respectively, has been putatively related to phenotypes as divergent as depression and antisocial personality disorder, respectively; both nuclei project to the amygdala ^46^. Recent data underscore the complexity of serotonin subsystems, revealing that even within the dorsal raphe there are subsystems that have distinct and at times opposing functions: both activate to reward but have opposing responses to aversion ^77^.

A limitation of our experiment is we did not measure serotonin (5-HT) directly: we measured plasma tryptophan levels following depletion, as tryptophan is the amino acid precursor of serotonin and ATD has been shown to produce transient reductions in central serotonin synthesis in humans ^11^. Whilst the validity of ATD as a method to manipulate central serotonin has been questioned ^78^, this position has been rebutted on the basis of considerable evidence ^9^. Consonant results between human studies employing ATD, and rodent experiments that induce profound serotonin loss using the neurotoxin 5,7-DHT, bolster the case that ATD reduces central serotonin. A prime example comes from studies of waiting impulsivity, which can be induced in humans following ATD ^49^, and in rats after serotonin depletion via 5,7-DHT ^79^. It is also important to note that the present task did not measure positive emotions. Whilst ATD is associated with evoking negative biases ^2^, consistent with our results, future work will be required to clarify whether positive emotions to social scenarios would be blunted or potentiated by ATD.

Our data from a novel test, that required drawing on autobiographical memories to mentally simulate cued social scenarios, demonstrate that there are important individual differences in the way serotonin influences how we react emotionally to social injustice. This should not come as a surprise given the intricacy of the serotonin systems and the complexities of human emotion and behaviour. Whilst serotonin depletion potentiated the magnitude of emotion non-specifically at the group level, personality traits played a critical role in shaping which distinctive types of emotions were affected. There was a qualitative dissociation in the way trait empathy, relative to psychopathy, amplified social emotion following serotonin depletion. Previous ATD studies on social cognition, by contrast, examined behavior rather than emotion and found changes in the magnitude but not the quality of effects ^5,6^. We propose that traits in conjunction with the memories our task evoked represent biological priors, which prime individuals to have different emotional reactions in the social world. Our data indicate serotonin would affect the gain. Given emotions are a prescription for action ^65^, it follows that our results could represent how serotonin impacts social behaviour via underlying emotional responses, positioned at the nexus of a social Pavlovian influence over action (Pavlovian action selection). When considering apparent paradoxes in the serotonin literature ^2,46^ and designing future studies, it is critical to note that the quality and magnitude of effects of a single serotonin manipulation can depend on personality. These data additionally inform the neurochemical basis of psychopathology associated with excessive emotions such as guilt and shame. Our findings on the interaction between the serotonin depleted state and personal attributes could help inform which individuals are particularly vulnerable to pathological emotional reactions, and who may be more amenable to serotonin-modulating treatments, with implications for psychiatric classification in frameworks such as the Research Domain Criteria ^80^.

## Acknowledgements

This research was funded by a Wellcome Trust Senior Investigator Award (104631/Z/14/Z) to T.W.R. B.J.S. receives funding from the National Institute for Health Research (NIHR) Cambridge Biomedical Research Centre (Mental Health Theme); the views expressed are those of the authors and not necessarily those of the NIHR or the Department of Health and Social Care. R.N.C.’s research is supported by the UK Medical Research Council (MC_PC_17213). J.W.K. is supported by a Gates Cambridge Scholarship. We would like to thank the staff at the NIHR/Wellcome Trust Clinical Research Facility at Addenbrooke’s Hospital, where the study was conducted, and Rachel Kyd of the Cambridge University Hospital Research & Development Office for assistance with study approval.

## Conflict of Interests statement

T.W.R. discloses consultancy with Cambridge Cognition, Greenfields Bioventures and Unilever; he receives research grants from Shionogi & Co and GlaxoSmithKline and royalties for CANTAB from Cambridge Cognition and editorial honoraria from Springer Verlag and Elsevier. B.J.S discloses consultancy with Cambridge Cognition, Greenfield BioVentures, and Cassava Sciences, and receives royalties for CANTAB from Cambridge Cognition. R.N.C. consults for Campden Instruments and receives royalties from Cambridge Enterprise, Routledge, and Cambridge University Press. J.W.K., F.E.A., R.Y, D.M.C., A.M.A-S., and A.P. declare no conflicts of interest.

