## SupplementaryMaterial for "Serotonin depletion amplifies distinct human social emotions as a function of individual differences in personality"

### Supplementary Material

### Methods

##### Further exclusion criteria

Further exclusion criteria: history of taking endocrine medication, use of St. John's Wort, pregnancy, regular consumption of over 38 units of alcohol per week, consumption of more than five cigarettes per day, use of cannabis more than once per month, use of other recreational drugs besides cannabis more than five times in the lifespan, cardiac or circulation problems, respiratory issues including asthma, gastrointestinal disorders, kidney disorders, thyroid problems, head injury, a bleeding disorder, and diabetes.

##### Individual characteristics

We assessed several personality traits and psychological symptoms primarily to ensure the placebo and depletion groups were matched: trait anxiety ($t_{(71)}$ = -0.872, p = .386) using the Spielberger Trait Anxiety Inventory (STAI; Spielberger et al., 1983); autistic characteristics ($t_{(71)}$ = -0.112 , p = .911) using the Adult Autism Spectrum Quotient (AQ; Baron-Cohen et al., 2001); Participants in each group, additionally, did not differ in their years of education ($t_{(71)}$ = 0.634, p = .528).

##### Principal Components Analysis Supplementary Information

Our principal components analysis (PCA) was performed with oblique rotation (direct oblimin). The Kaiser-Meyer-Olkin (KMO) measure verified the sampling adequacy for the analysis, KMO = .719 ("good" according to Hutcheson & Sofroniou, 1999). Bartlett's test of sphericity indicated correlations between items were sufficiently large for PCA ($\chi^2$ (120) = 700.731, p = 7.9941 x $10^{-83}$). We obtained eigenvalues for each component in the data and extracted components with eigenvalues over Kaiser's criterion of one. This resulted in 4 components, which explained 72.354% of the variance. The scree plot showed an inflexion that justified retaining these 4 components, which were then used in the remainder of the analysis. The pattern matrix (Supplementary Table 1) shows the factor loadings after rotation.

#### Figures and Tables

**Supplementary Figure 1.** Effects of ATD on emotion: serotonin depletion enhanced emotion non-specifically overall (see main text). Each bar represents the average emotion ratings per group, collapsed across agency and intentionality. Error bars indicate 1 standard error. Asterisk indicates main effect, $p < .05$.

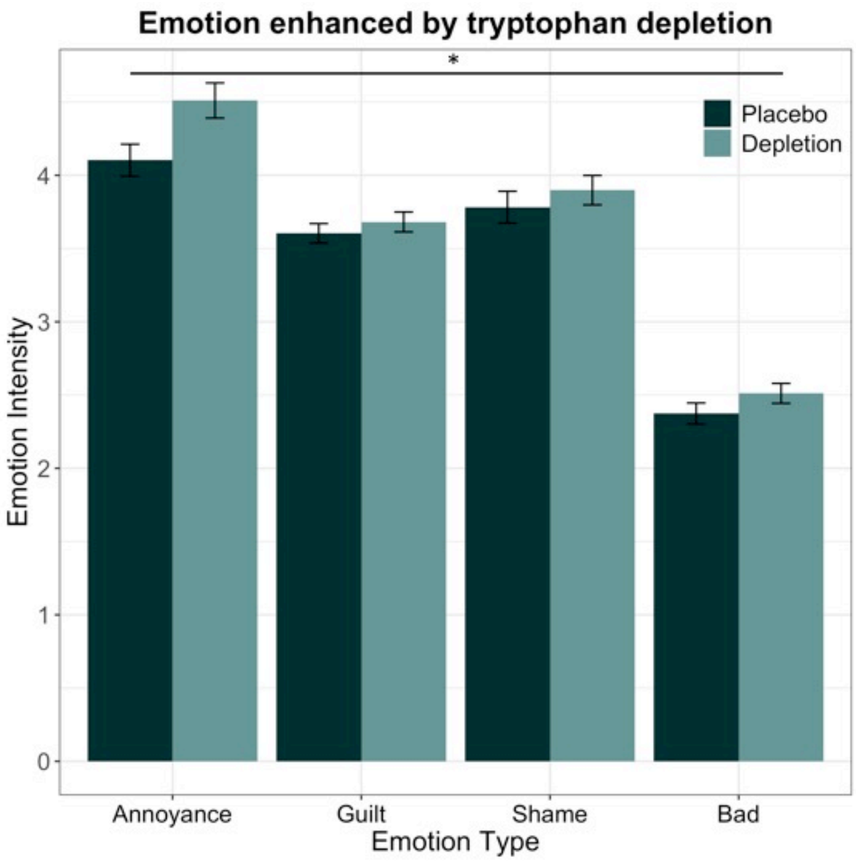

**Supplementary Figure 2.** ANOVA on four principal components from principal components analysis. Measures contained in components 2 and 4 were significantly increased by ATD (acute tryptophan depletion). Error bars indicate 1 standard error. # indicates interaction at $p < .05$; asterisk indicates significant follow-up t-test at $p < .05$.

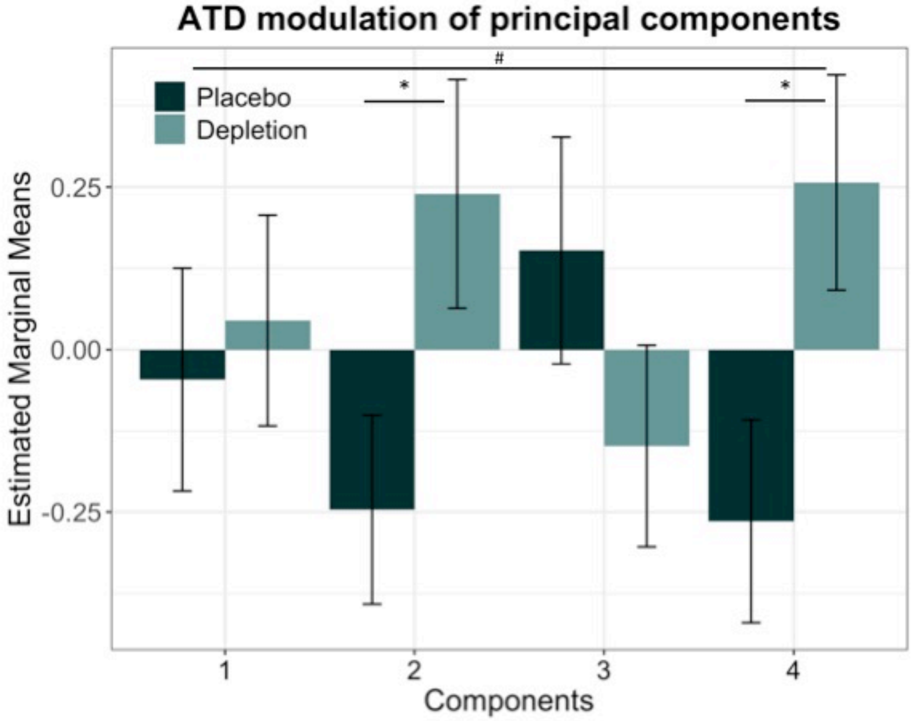

**Supplementary Table 1.** Principal Components Pattern Matrix. Rotation Method: Oblimin with Kaiser Normalisation. Rotation converged in 19 iterations. Intended = intentional harm. Unintended = unintentional harm.

| | **Component** | | | |
|---|---|---|---|---|
| **Raw Variable** | **1** | **2** | **3** | **4** |
| Annoyed-unintended-victim | .819 | | | |
| Annoyed-intended-victim | .791 | .317 | | |
| Bad-unintended-victim | -.736 | | | |
| Bad-unintended-agent | -.617 | | -.351 | |
| Bad-intended-victim | -.604 | | | |
| Guilty-unintended-agent | .587 | | .403 | |
| Annoyed-unintended-agent | | .921 | | |
| Annoyed-intended-agent | | .897 | | |
| Ashamed-intended-agent | | | .923 | |
| Guilty-intended-agent | | | .922 | |
| Bad-intended-agent | | | -.892 | |
| Ashamed-unintended-agent | .396 | | .455 | .417 |
| Ashamed-unintended-victim | | | | .830 |
| Ashamed-intended-victim | | | | .750 |
| Guilty-unintended-victim | | .408 | | .541 |
| Guilty-intended-victim | | .504 | | .521 |

**Supplementary Table 2.**

| | **Guilt** | **Shame** | **Bad** |
|---|---|---|---|
| **Annoyance** | r = .481, p = $1.7 \times 10^{-5}$ | r = .403, p = $4.08 \times 10^{-4}$ | r = -.145, p = .220 |
| **Guilt** | | r = .779, p = $5.16 \times 10^{-16}$ | r = -.527, p = $2.0 \times 10^{-6}$ |
| **Shame** | | | r = -.451, p = $6.2 \times 10^{-5}$ |
